# Prenatal methadone exposure leads to long-term memory impairments and disruptions of dentate granule cell function in a sex-dependent manner

**DOI:** 10.1101/2022.03.22.485361

**Authors:** Meredith E. Gamble, Rhea Marfatia, Marvin R. Diaz

## Abstract

Prenatal opioid exposures lead to extensive cognitive and emotion-regulation problems in children, persisting at least through school-age. Methadone, an opioid typically used for the treatment of opioid use disorder, has been approved for use in pregnant women for several decades. Importantly, however, the impacts of prenatal methadone exposure (PME), particularly on offspring as they progress into adulthood, has not been extensively examined. In recent years, children and young animal models have shown cognitive deficits related to PME, including evidence of hippocampal dysfunction. The present work aims to examine the persistent nature of these deficits, as well as determine how they may differ by sex. Pregnant Sprague Dawley rats either received subcutaneous methadone or water injections twice daily from gestational days 3-20 or were left undisturbed. Following postnatal day 70, male and female offspring were behaviorally tested for impairments in recognition memory using the Novel Object Recognition task and working spatial memory through Spontaneous Alternation. Additionally, using whole-cell patch-clamp electrophysiology, hippocampal dentate granule cell function was examined in adult offspring. Results indicate that methadone-exposed females showed decreased excitability and increased inhibition of these dentate granule cells, while males did not. These findings were accompanied by impairments in female working spatial memory and impaired recognition memory of both sexes. Overall, this work supports the continued investigation of the long-term effects of PME on adult male and female learning and memory, as well as promotes further exploration of adult hippocampal function as a neural mechanism impacted by this exposure.

## 1. Introduction

The rates of opioid use disorder (OUD) in the United States have drastically escalated over the past two decades, including opioid misuse among pregnant women. The rates of maternal opioid misuse at the time of delivery has quadrupled during this time, and the number of infants diagnosed with neonatal opioid withdrawal syndrome (NOWS) has risen from 1.5 to 6.5 cases for every thousand hospital births^1^. Importantly, OUD treatments, such as methadone and buprenorphine, have been approved for use in pregnant women for several decades^2,3^, with early reports showing methadone produces better outcomes for the infant compared to other misused opioids^2,4,5^. However, as methadone continues to be the standard of care, more recent data has begun to uncover deficits in children beyond infancy following exposure to methadone *in utero*.

More specifically, prenatal methadone exposure (PME) significantly reduces IQ scores in children aged 3 to 6 years^6^ and increases the rates of reading and math delays at age 9.5 years^7^. These findings have been supported by preclinical studies showing PME leads to impairments in recognition and working memory of adolescent rats^8,9^ and long-term spatial memory of young adult rats^10^. These findings indicate that while PME may result in more favorable neonatal outcomes, it may produce similar deficits as shown with prenatal exposures to other opioids, such as oxycodone and morphine, later on. These include decreased retention and spatial memory in juvenile rodents^11,12^ and impairments in working, spatial, and retention memory during adolescence^13,14^ following prenatal morphine exposures. In adult offspring, data have shown impaired spatial memory following prenatal l-alpha-acetylmethadol (LAAM) exposure^15^ and prenatal oxycodone exposure^16^, and impaired reference and recognition memory following prenatal buprenorphine exposure^10^. These behavioral deficits have been attributed to dysfunction of several neural mechanisms, notably hippocampal and prefrontal cortex dysfunction in young rodents, including decreased neurogenesis and neuroplasticity in these regions^11,13,14,17,18^.

While these studies are essential in continuing to evaluate the dangers of opioid use during pregnancy, including prescribed methadone, the persistent nature of PME-related effects throughout the lifetime remains unknown. Therefore, it is challenging to predict how this exposure may affect the development of exposed children as they continue to age. It is unclear whether PME-induced deficits persist into adulthood or emerge later in life. This is particularly important as the surging rates of OUD have initiated over the past two decades, with the children exposed during this time not yet reaching adulthood.

The present study aimed to begin characterizing adult male and female memory impairments following PME, including assessment of granule cell function within the dentate gyrus (DG) of the hippocampus. We selected this region as one that may be disrupted by this exposure, as most animal behavioral work described above relate to deficits in DG-dependent tasks^19–23^.

## 2. Methods

### 2.1. Animals

Male and female Sprague-Dawley rats (N = 172, from approximately 50 litters) were bred in-house, with breeding pairs originating from Envigo (Indianapolis, IN). At weaning [postnatal (P) day 21], same-sex littermates were housed two to three per cage and randomly assigned to an experimental condition. Rats continued to receive food (5L0D PicoLab Laboratory Rodent Diet) and water *ad libitum* and were maintained on a 12-hr light-dark cycle (lights on at 07:00). All rats were treated following guidelines for animal care under protocols approved by Binghamton University Institutional Animal Care and use Committee.

### 2.2. Breeding and Prenatal Methadone Exposure (PME)

The PME used was adapted from previous studies^8,24^. For breeding, two naïve females were placed in a cage with one naïve male for up to four days. Females were checked every morning for pregnancy via vaginal smears, with the first day of detectable sperm designated gestational day (G)1^25^. Once pregnancy was confirmed, females were single-housed and randomly assigned to an exposure condition (methadone, water, or naïve) and continued to receive food (Purina Lab Diet 5008C33) and water *ad libitum*. PME began on G3, with dams receiving twice daily subcutaneous injections, twelve hours apart (07:00 and 19:00), of 5 mg/kg of either methadone dissolved in sterile water or sterile water alone. Dams continued to receive twice daily injections of 7 mg/kg on G4-20. Due to gnawing and pica side effects of the methadone exposure and to prevent ingestion of bedding and nesting material, both the water- and methadone-exposed dams were placed in a cage with no bedding, but continued access to food and water, for three hours following each injection. During this time, animals were closely monitored for side effects such as catalepsy and comatose behavior, in which case they would be placed on a heating pad and received artificial tears until recovered. After three hours, they were returned to their homecage until the next injection. Skin irritation was observed in some methadone-injected females (this was never observed in any water-injected females) and was treated with 1% silver sulfadiazine topical cream until resolved. Following G20, dams were left undisturbed to give birth. Daily weights were taken for all dams throughout the exposure. Those assigned to the naive group were left undisturbed throughout gestation except for the daily weights, in order to control for procedural stress (twice daily injections and temporary rehousing in bedding-free cages). After giving birth, dams were left undisturbed with their pups for two days. On P2, pups were counted, weighed, and culled to twelve (equal sex ratio when possible) and returned to the mother until weaning (P21). Throughout this time, pups were also weighed on P7 and P12, and were monitored to determine at what postnatal day they opened their eyes, for assessment of developmental delay. Following weaning, offspring were left to age until adulthood testing (P70+) (**Fig.1**).

**Figure 1.**
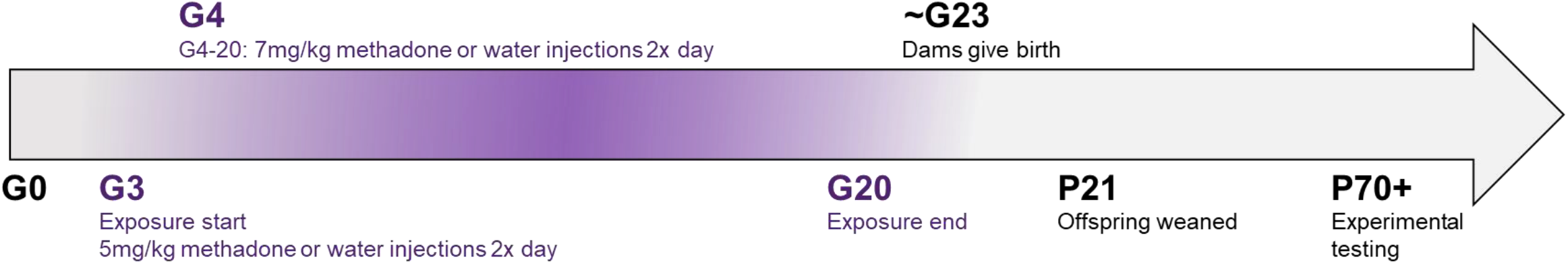
Prenatal methadone exposure (PME) timeline. Pregnancy was confirmed on gestational day (G) 1, and PME began on G3 with 5mg/kg of either methadone or water. Exposures continued at 7mg/kg G4-20, after which they were left undisturbed until they gave birth (~G23). Offspring were weaned on postnatal day (P) 21, and then left to age until testing began in adulthood (P70+).

### 2.3. Whole-cell patch-clamp electrophysiology

Adult offspring (P70-120; n = 5-7 per group, per sex across 2-5 litters per group) were briefly sedated with 3% isoflurane and quickly decapitated. Brains were rapidly removed and immersed in ice cold oxygenated (95% O_2_, 5% CO_2_) sucrose artificial cerebrospinal fluid (ACSF) cutting solution containing (in mM): sucrose (220), KCl (2), NaH_2_PO_4_ (1.3), NaHCO_3_ (26), glucose (10), MgSO_4_ (12), CaCl_2_ (0.2), and ketamine (0.43). 300 μm brain slices containing the dorsal hippocampus were made using a Vibratome (Leica Microsystems, Bannockburn, IL, USA). Slices were incubated in oxygenated ACSF containing (in mM): NaCl (125), KCl (2), NaH_2_PO_4_ (1.3), NaHCO_3_ (26), glucose (10), CaCl_2_ (2), MgSO_4_ (1), and ascorbic acid (0.4), and allowed to recover for at least 40 min at 34°C before recording. Slices remained at 34°C, and all experiments were performed within 5 hours of slice preparation.

Following incubation, slices were transferred to a recording chamber in which oxygenated ACSF was heated to 32°C and superfused over the slice at 3 mL/min. Dentate granule cells (DGC) from the dorsal hippocampus were identified based on regional appearance, morphology, and capacitance (<150 pF), with a threshold for acceptable access resistance as < 30 MΩ. Recordings from DGC were visualized using infrared-differential interreference contrast microscopy (Olympus America, Center Valley, PA). Spontaneous excitatory postsynaptic current (sEPSC) and intrinsic excitability recordings were collected with patch pipettes filled with a K-gluconate internal solution containing (in mM): K-Gluconate (120), KCl (15), EGTA (0.1), HEPES (10), MgCl_2_ (4), MgATP (4), Na_3_GTP (0.3), phosphocreatine (7), with a pH of 7.3 and osmolarity of 295-305 mOsm. sEPSC recordings were collected following pharmacological blockade of GABA_A_ receptors using 10 mM gabazine. For intrinsic excitability, resting membrane potential (RMP) and depolarization steps, recordings took place in a bath of either 1) ACSF, 2) 10 mM gabazine (to block GABA_A_ receptors), and 3) 10 mM gabazine, 1 mM kynurenic acid (KA), and 50 μM DL-APV (to block GABA_A_, AMPA, and NMDA receptors, respectively). In a separate subset of cells, GABA_A_ receptor-mediated spontaneous inhibitory postsynaptic current (sIPSC) recordings were collected using patch pipettes filled with a KCl internal solution containing (in mM): KCl (135), HEPES (10), MgCl_2_ (2), EGTA (0.5), Mg-ATP (5), Na-GTP (1) and QX314-Cl (1), with a pH of 7.25 and osmolarity of 280-290 mOsm. sIPSCs were recorded following pharmacological blockade of AMPA and NMDA glutamate receptors using 1 mM KA and 50 μM DL-APV. Data were acquired with MultiClamp 700B (Molecular Devices, Sunnyvale, CA) at 10 kHz, filtered at 1 kHz, and stored for later analysis using pClamp software (Molecular Devices).

Upon rupturing the cell membrane, cells were allowed to dialyze for at least 5 min before recording began. For intrinsic excitability, recordings began with RMP and depolarization steps using current-clamp configuration in a bath of ACSF. For depolarization steps, cells were held at −70 mV to account for any individual differences in their native RMP. Steps were recorded beginning at −50 pA in successive increasing steps of 15 pA with a 500 ms duration. Access resistance was monitored throughout each recording, and neurons in which the access resistance changed > 20% were not included in analysis.

### 2.4. Novel Object Recognition

A separate group of adult offspring (n = 8-10 per group, per sex across 6-10 litters per group) were tested for PME-induced impairments in recognition memory using the Novel Object Recognition (NOR) task. Testing included three days: habituation, familiarization, and test day. Animals were handled for 2 days prior to the habituation day. For habituation, animals were placed in the center of an empty 40 cm x 40 cm plexiglass arena and allowed to explore for 10 minutes. Twenty-four hours after habituation, animals were placed in the same arena with two identical objects and allowed to explore the objects for 10 minutes. On test day, one of the identical objects was replaced with a novel one. Animals were again placed in the arena with these two objects and allowed to explore for 5 minutes. Novel and familiar objects were either Lego statues or wooden blocks, counterbalanced between animals (substantial exploration of these objects was determined previously in pilot experiments). Video recordings were used for data scoring and analyses by an experimenter blind to exposure conditions. Videos were scored for duration of time spent exploring the objects (sniffing and direct contact with the object) during the first three minutes, and a discrimination ratio was calculated for the test.

### 2.5. Spontaneous Alternation

We also examined working spatial memory in a separate group of animals (n = 8 per group, per sex across 4 litters per group) using the Spontaneous Alternation task. Following seven days of handling, animals were placed one at a time in an opaque habituation cage for 30 minutes, to adjust to the room lighting and environment. Following habituation, animals were placed in the center of a plus-maze for eighteen minutes surrounded by spatial cues on the walls of the room. Experimenters, blind to the animals’ conditions, stayed in the room and recorded each arm entry (all four paws in the arm) throughout the testing period. An alternation is defined as entry into each of the four arms in a successive sequence, without any repeats (i.e. 1,3,2,4). Percent alternation scores were calculated as ((number of alternations/ (total number of arm entries – 3)) *100), as previously described^26,27^

### 2.6. Drugs and Chemicals

Chemicals were purchased from Sigma-Aldrich (St. Louis, MO) unless otherwise noted. Kynurenic acid, DL-APV, and QX314-Cl were purchased from Tocris/R&D Systems (Bristol, UK). Methadone hydrochloride was acquired through the NIH Drug Supply Program and purchased from Sigma-Aldrich.

### 2.7. Statistics

Statistical analyses for all data were completed using Prism 8 (GraphPad, San Diego, CA). All data were first tested for normality (Kolmogorov-Smirnov test), and nonparametric tests were used when data were not normally distributed. Dam weights were analyzed first with a one-way ANOVA to rule-out any significant differences at the start of exposure (G3), then gestation weights across exposure and pup weights were analyzed with a mixed-effects model (exposure x gestational or postnatal day) using the Geisser-Greenhouse correction for sphericity. Tukey’s multiple comparison post-hoc test was used for follow-up on any significant effects. Eye-opening data were analyzed using a two-way repeated-measures ANOVA, and percent female pups in each litter were analyzed using a one-sample t-test compared to chance (0.5) and a one-way ANOVA between exposure groups.

Electrophysiological data were analyzed using MiniAnalysis (Synaptosoft Inc.) and Clampfit (Molecular Devices). For electrophysiology excitability analyses, data included cells that survived through all three drug washes. A one-way ANOVA was run for each drug wash experiment independently. If data did not pass the normality test, a non-parametric (Kruskal-Wallis test) was used instead. A two-way repeated-measures ANOVA was conducted to determine any differences in the current-action potential (AP) curves for each washed-on drug. One-way ANOVAs were run for exposure differences in sIPSC and sEPSC frequency, amplitude, event decay, AP time and threshold measures, as described below.

Due to the sex-dependent findings in our electrophysiology results, as well as the substantial literature on sex differences in performance of rodents in learning and memory tasks^28–30^, we have separated analyses of both behavioral tasks by sex. NOR data were analyzed using a one-sample t-test comparing mean discrimination ratio values to 0.5 (chance) or 0 (percent change from naïve controls). Data were analyzed with a one-way ANOVA test for differences between exposure groups. Female NOR discrimination ratio data did not pass the test for normality and were run using Wilcoxon signed-rank test. Spontaneous alternation data were analyzed using a one-way ANOVA. Tukey’s multiple comparison post-hoc test was used for follow-up on any significant effects.

## 3. Results

### 3.1 Characterization of PME Model

To characterize the impact of this PME model on early growth and development, we took daily gestation weights of the dams, weights of the pups following birth, and identified the postnatal day the pups opened their eyes, as a measure of a typical development milestone. Methadone-exposed dams showed a significant decrease in body weight during the exposure period (**Fig. 2A;** *F*(2,43)=5.292, *p*=0.009), with Tukey’s post hoc tests revealing emerging delays in weight gain starting on G7 (*p*=0.029) continuing through G20 (*p*<0.0001) compared to naïve dams. Water-exposed dams did not show any significant differences in weight compared to naïve dams. There were no significant differences of dam weight at the start of exposure on G3 (*F*(2,43)=0.693, *p*=0.506).

**Figure 2.**
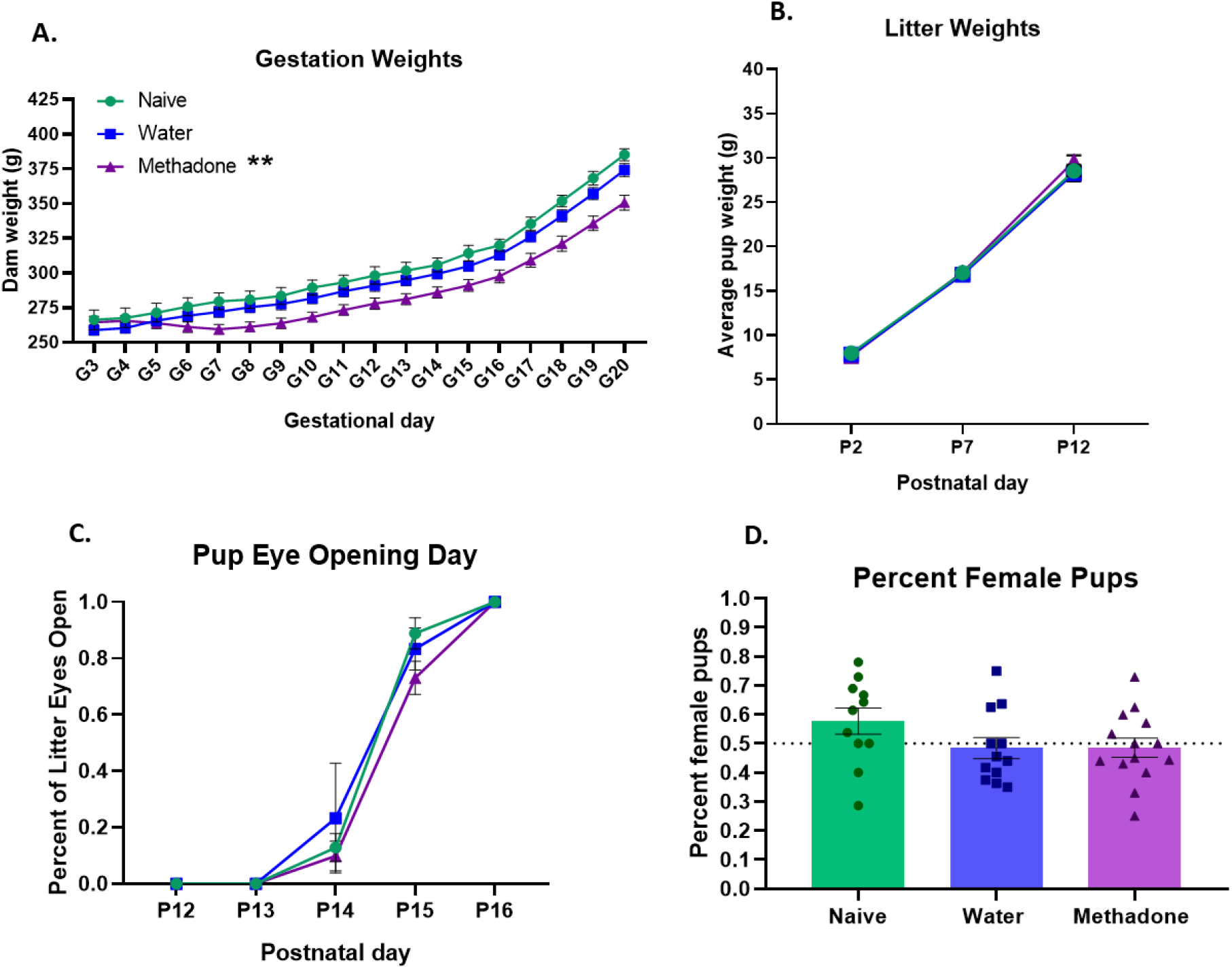
PME litter characterization. Methadone-exposed dams showed a significant reduction in weight throughout the exposure period, with no significant differences at the start of exposure (gestational day [G] 3) (A). However, offspring did not show any significant differences in developmental weight (B) or the postnatal day (P) they opened their eyes (C). Finally, there were no significant differences in pup sex ratios as a result of prenatal methadone or water exposures (D), with individual data points representing litters; error bars represent SEM. ** = p < 0.01 main effect exposure.

Following birth, pups were weighed on P2, 7, and 12 with no significant differences in weight as a result of exposure condition (**Fig. 2B**; *F*(1.94, 88.27)=8.56, *p*=0.426) and no significant day x exposure interaction (*F*(4, 91)=0.888, *p*=0.475). Pups were also monitored to see at what postnatal day they opened their eyes, with no significant differences across the three exposure conditions (**Fig. 2C**; *F*(2,17)=0.833, *p*=0.452). Finally, the sex ratio of surviving pups was also analyzed, revealing no significant differences in the percent of female pups among the three exposure conditions when compared to 50% (**Fig. 2D**; naïve: *t*(10)=1.723, *p*=0.116; water: *t*(11)=0.435, *p*=0.672; methadone: *t*(13)=0.427, *p*=0.676) or across the three groups (*F*(2,43)=1.883, *p*=0.168).

### 3.2 PME alters dentate granule cell excitability in a sex-dependent manner

Both the RMP and the rheobase of each cell were recorded as measures of intrinsic excitability with 1) all synaptic input intact (ACSF), 2) only GABA transmission blocked (+gabazine), and 3) GABA and glutamate transmission blocked (+gabazine, KA, APV). For females, results indicate no significant differences across exposure groups for RMP (**Fig. 3A**; ACSF: *F*(2,28)=1.070, *p*=0.357; gabazine: *H*(2)=1.815, *p=* 0.404; gabazine, KA, APV: *H*(2)=3.484, *p=*0.175).

**Figure 3.**
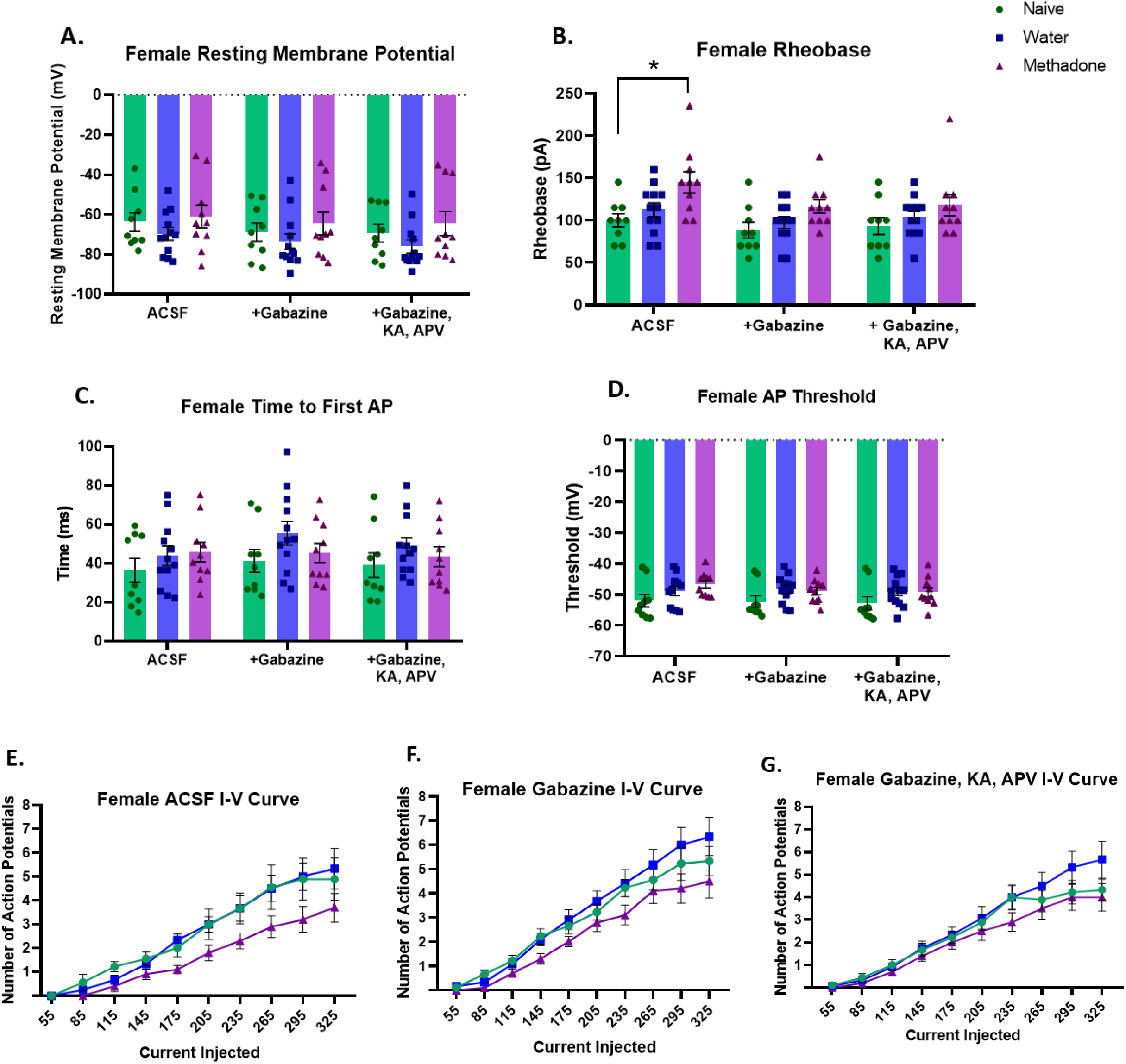
PME leads to decreased excitability of dentate granule cells in adult female offspring. Females did not show any significant changes to resting membrane potential in any of the three drug conditions (artificial cerebrospinal fluid [ACSF], gabazine, or gabazine, kynurenic acid and APV-5 [KA, APV]) (A); however, PME females showed a significantly higher rheobase in the ACSF condition, indicating decreased excitability of these cells compared to controls, but this effect did not persist when blocking synaptic activity in the other two drug conditions (B). Females did not show any significant changes to either time to first AP (C) or threshold (D). When examining frequency of action potential firing with depolarization steps, females did not show any significant differences in the number of action potentials fired at each depolarization step for ACSF (E), gabazine (F), or gabazine, KA, and APV (G). Data points represent individual cells with up to three cells per animal; error bars represent SEM. * p < 0.05 compared to naïve.

For the rheobase, females showed a significant difference in the ACSF bath (*F*(2,28)=5.289, *p*=0.011), with post hoc analysis revealing significant increases in the methadone animals compared to naive (*p*=0.012), with no significant difference between naïve and water-exposed offspring (*p*=0.645). In contrast, there were no significant rheobase differences when blocking GABA activity (*F*(2,28)=3.026, *p*=0.065), or when all synaptic activity was blocked (*H*(2)=2.231, p=0.328) (**Fig. 3B**).

Additional AP kinetics were analyzed, including time to first AP following current injection at rheobase and the membrane potential at the start of the first AP (threshold) for each cell. Females showed no significant differences in time to first AP between the exposure groups (ACSF: *F*(2,28)=0.803, *p*=0.458; gabazine: *F*(2,28)=1.661, *p*=0.208; gabazine, KA, APV: *F*(2,28)=0.876, *p*=0.428; **Fig. 3C**). Additionally, females showed no significant threshold difference across the exposure groups (ACSF: *F*(2,28)=2.498, *p*=0.100; gabazine: *H*(2)=4.381, *p*=0.112; gabazine, KA, APV: *H*(2)=4.367, *p*=0.113; **Fig. 3D**).

Next, the number of APs induced with increasing depolarization steps were recorded to create a current-AP curve. There were no significant differences among the three exposure conditions for the number of APs fired under any of the three drug conditions. For ACSF, results indicated a significant increase in AP frequency across current injections, as expected, but no significant effect of exposure (*F*(2,28) 2.544, *p*=0.097) and no significant interaction (*F*(18,252)=1.384, *p*=0.139). Results were similar for the gabazine (exposure: *F*(2,28)=2.084, *p*=0.143; interaction: *F*(18,252)=1.224, *p*=0.242) and the gabazine, KA, and APV condition (exposure: *F*(2,28)=1.186, *p*=0.320; interaction: *F*(18,252)=1.253, *p*=0.219) (**Fig. 3E-G**).

For males, results indicated no significant differences in any of the major excitability measures. There were no significant differences in RMP (ACSF: *p*=0.394; Gabazine: *p*=0.213; Gabazine, KA, APV: *p*=0.457) (**Fig. 4A**) or rheobase for any of the exposure or drug wash conditions (ACSF: *p*=0.646; Gabazine: *p*=0.875; Gabazine, KA, APV: *p*=0.564) (**Fig. 4B**).

**Figure 4.**
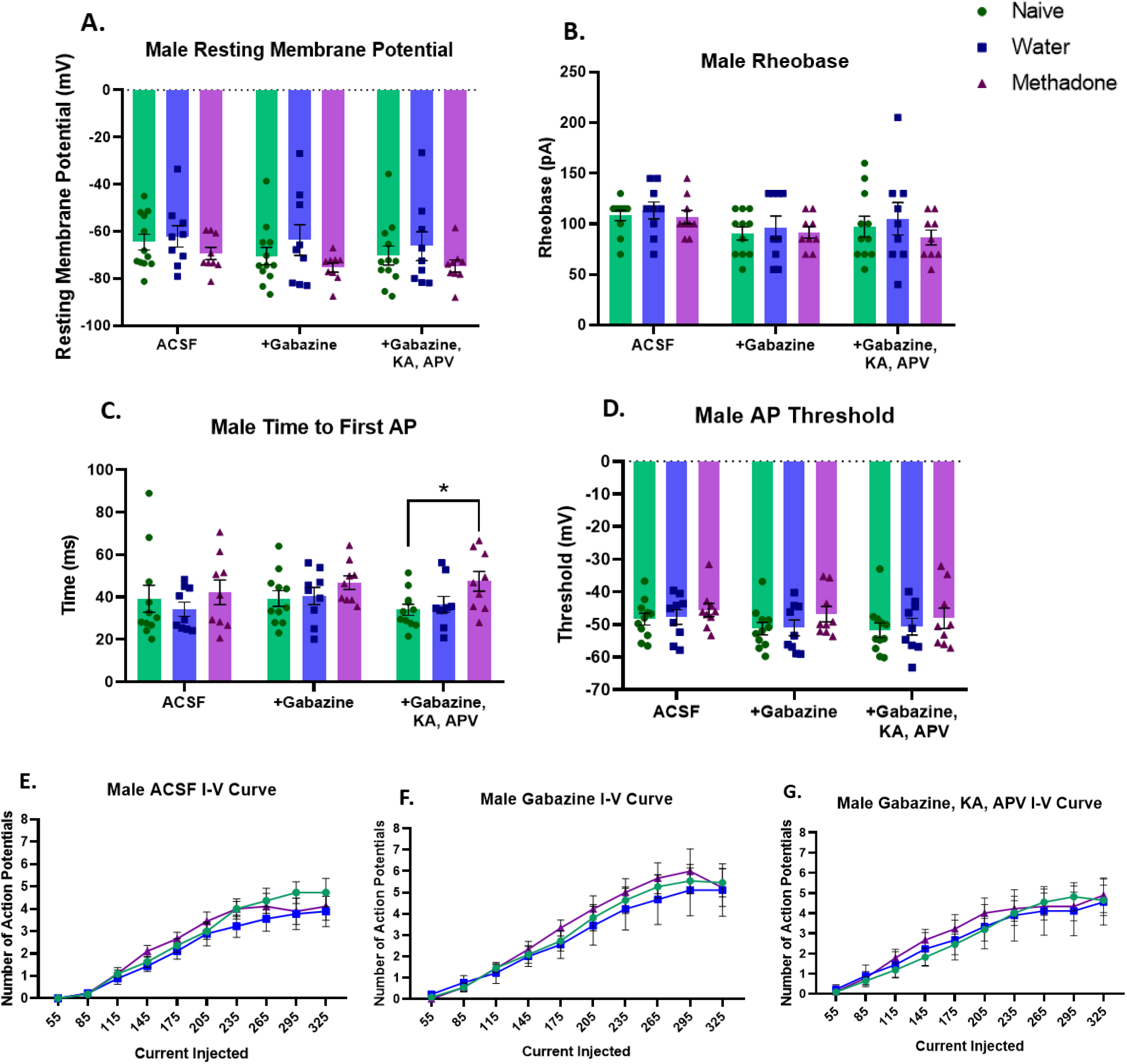
PME does not significantly alter male dentate granule cell excitability. Males did not show any significant changes to resting membrane potential in any of the three drug conditions (artificial cerebrospinal fluid [ACSF], gabazine, or gabazine, kynurenic acid and APV-5 [KA, APV]) (A), nor any significant changes to rheobase across any of the drug conditions (B). PME males did show a significant increase in time to first action potential when blocking synaptic activity (+gabazine, KA, APV), but not in the other drug conditions (C), and no changes to AP threshold (D). When examining frequency of action potential firing with depolarization steps, males did not show any significant differences in the number of action potentials fired at each depolarization step for ACSF (E), gabazine (F), or gabazine, KA, and APV (G). Data points represent individual cells with up to three cells per animal; error bars represent SEM. * p < 0.05 compared to naïve.

For time to first AP, males showed no significant differences in the ACSF bath (*H*(2)=2.258, *p*=0.533) or when blocking GABA transmission (*F*(2,26)=1.133, *p*=0.337); however, when blocking both glutamate and GABA activity, there was a significant difference in the time to AP (*F*(2,26)=3.754, *p*=0.037), with post-hoc test revealing methadone cells take longer to fire compared to naïve (*p*=0.043) (**Fig. 4C**). Finally, males showed no significant threshold differences across exposures (ACSF: *F*(2,26)=0.538, *p*=0.590; gabazine: *H*(2)=2.276, *p*=0.321; gabazine, KA, APV: *F*(2,26)=0.504, *p*=0.610; **Fig. 4D**).

There were also no significant effects for the male current-AP curves, indicating no exposure-related effects on frequency of AP firing (**Fig. 4E-G**). For ACSF, results indicated a significant increase across current injections, as expected, but no significant effect of exposure (*F*(2,26)=0.585, *p*=0.564) and no significant interaction (*F*(18,234)=0.747, *p*=0.760). Results were similar for the gabazine (exposure: *F*(2,26)=0.172, *p*=0.843; interaction: *F*(18,234)=0.357, *p*=0.993) and the gabazine, KA, and APV condition (exposure: *F*(2,26)=0.083, *p*=0.920; interaction: *F*(18,234)=0.358, *p*=0.993).

### 3.3 PME does not significantly alter glutamate transmission

Given that synaptic activity contributed to some differences in DGC excitability, we assessed glutamate neurotransmission onto DGC and examined both the frequency and amplitude of sEPSCs, as well as measures of event decay time (tau). Females showed no significant differences in sEPSC frequency or amplitude across exposure groups (frequency: *F*(2,26)=2.890, *p*=0.073; amplitude: *F*(2,25)=0.582, *p*=0.567; **Fig. 5A-C**). Similarly, males did not show any significant difference in sEPSC frequency or amplitude (frequency: *H*(2)=1.017, *p*=0.602; amplitude: *F*(2,24)=1.437, *p*=0.257; **Fig. 5D-F**). Additionally, neither sex showed any significant differences in decay time of sEPSCs (female tau 1: *F*(2,23)=0.911, *p*=0.416; female tau 2: *F*(2,23)=0.456, *p*=0.639; male tau 1: *H*(2)=0.831, *p*=0.660; male tau 2: *H*(2)=0.615, *p*=0.735; **Supplementary Fig. 1**).

**Figure 5.**
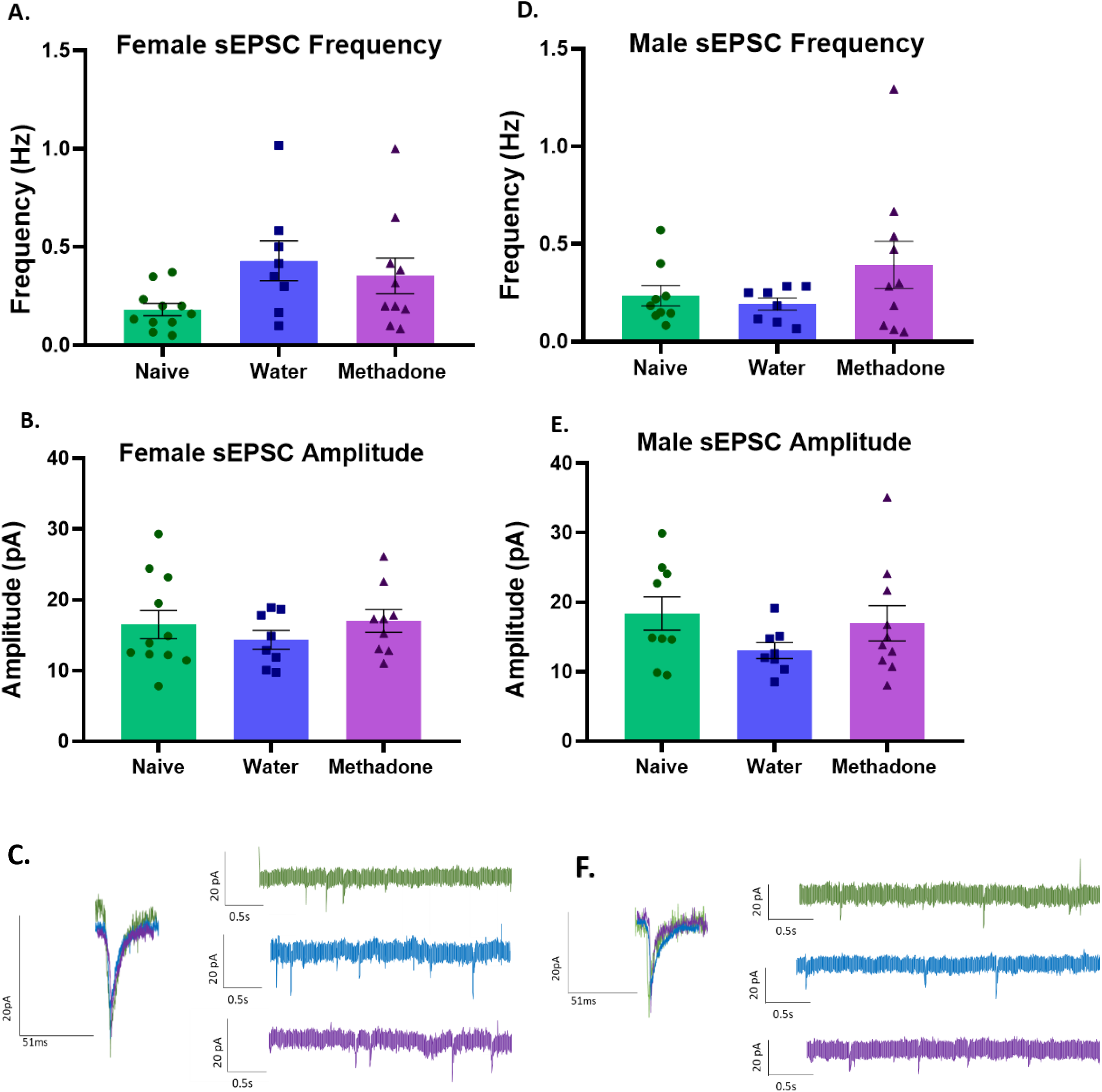
PME does not alter glutamate transmission. Females did not show any significant differences for sEPSC frequency (A) or amplitude (B), with representative traces shown (C), indicating no differences in glutamate transmission across the three exposure groups. Males also did not show any significant differences in sEPSC frequency (D) or amplitude (E), with representative traces shown (F). Individual data points represent cells, with up to three cells per animal; error bars represent SEM.

### 3.4 PME increases female, but not male, dentate granule cell inhibition

We also assessed GABAergic neurotransmission onto DGC. For females, results revealed that sIPSC frequency was not different (*F*(2,27)=3.302, *p*=0.052), yet sIPSC amplitude was significantly different (*F*(2,27)=3.448, *p*=0.046), with post hoc analysis revealing this effect is driven by an increase in amplitude of methadone-exposed females compared to naïve (*p*=0.037) (**Fig. 6A-C**). However, males did not show a significant difference in sIPSC frequency or amplitude (frequency: *H*(2)=0.365, *p*=0.833; amplitude: *F*(2,24)=0.131, *p*=0.878; **Fig. 6D-F**).

**Figure 6.**
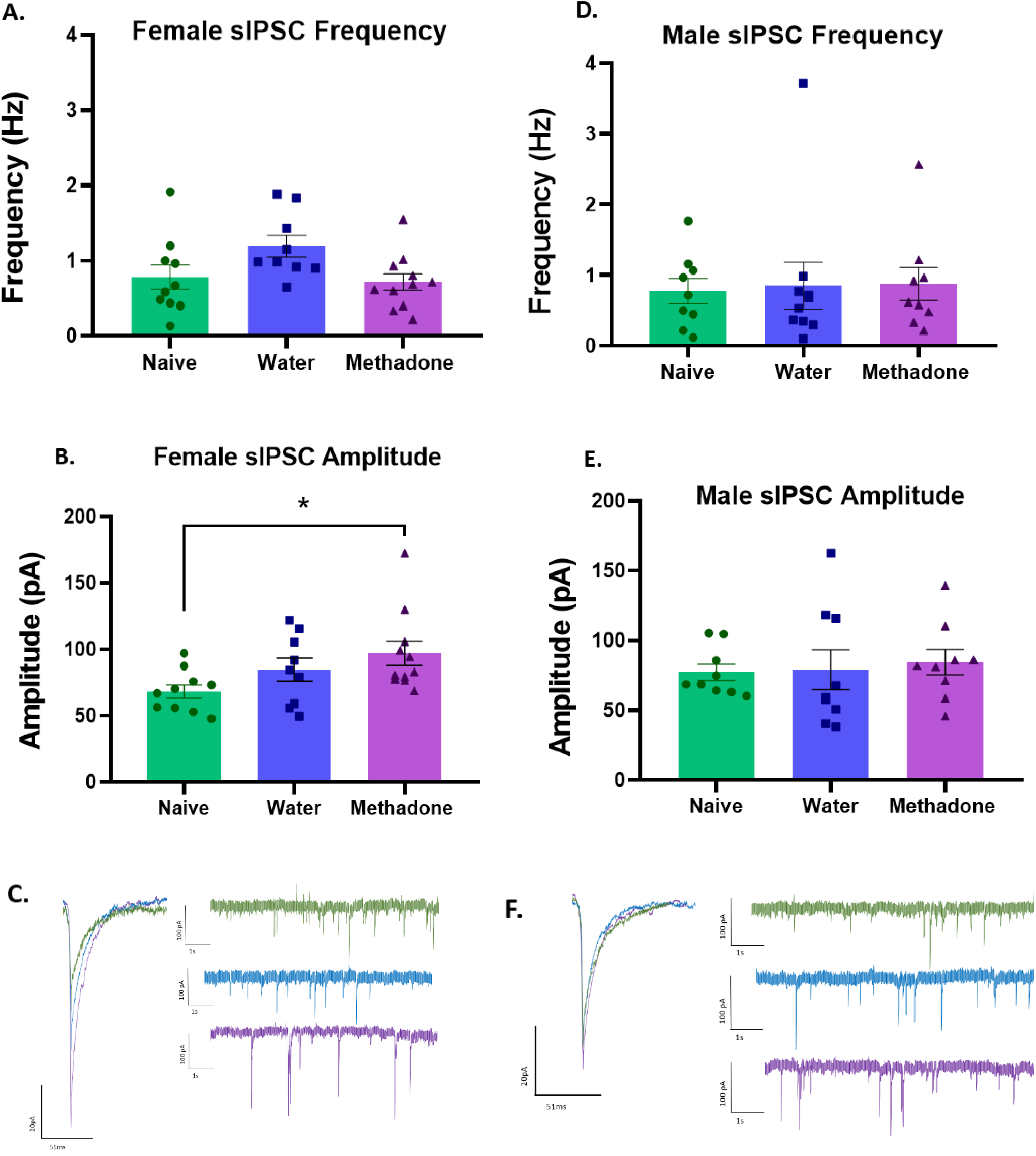
PME increases dentate granule cell inhibition only in females. Females did not show any significant differences in sIPSC frequency (A); however, PME females showed increased sIPSC amplitude (B), with representative traces shown (C). Males did not show any significant difference in sIPSC frequency (D) or amplitude (E), with representative traces shown. Individual data points represent cells, with up to three cells per animal. Error bars represent SEM; * p < 0.05 compared to naïve.

Neither sex showed significant difference in sIPSC event decay time (female tau 1: *H*(2)=0.989, *p*=0.610; female tau 2: *F*(2,22)=0.404, *p*=0.672; male tau 1: *F*(2, 17)=0.455, *p*=0.642; male tau 2: *F*(2,17)=0.339, *p*=0.717; **Supplementary Fig. 1**).

### 3.5 PME impairs performance on Novel Object Recognition task

Adult offspring were tested using the NOR task as a measure of recognition memory. For females, methadone-exposed data were not normally distributed, so a Wilcoxon Signed Rank Test was used. PME females showed significant reductions in discrimination ratios compared to chance (theoretical mean = 0.5) (*Mdn* = 0.459, *p*=0.039), while naïve and water-exposed offspring did not differ from chance (naïve: *Mdn* = 0.644, *p*=0.148; water: *Mdn* = 0.594, *p*=0.275). Additionally, females showed a significant difference in discrimination ratios across exposures (*H*(2)=7.115*, p*=0.029), with Dunn’s post-hoc test revealing reduced scores in PME females compared to naïve (*p*=0.027) (**Fig. 7A**). Similarly, PME males showed significant reductions compared to chance (*t*(7)=3.541, *M*=0.378, *p*=0.009), while naïve and water-exposed did not significantly differ from chance (naïve: *t*(7)=1.21, *M*=0.569, *p*=0.266; water: *t*(9)=0.794, *M*=0.555, *p*=0. 448). However, males did not show a significant difference across exposure groups (*F*(2,23)=3.190, *p*=0.060)(**Fig. 7B**).

**Figure 7.**
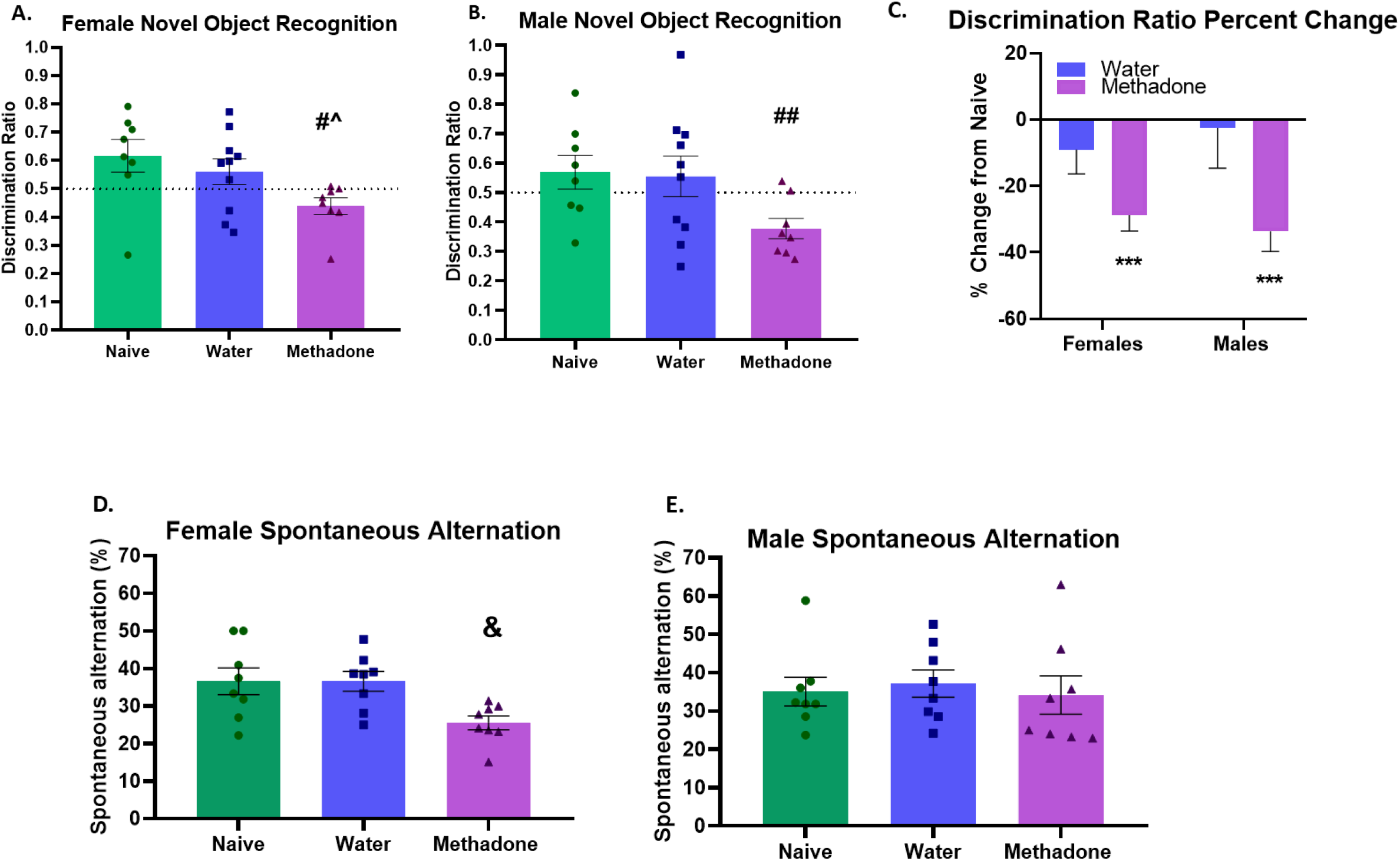
PME impairs female and male recognition memory and female working spatial memory in adult offspring. Both PME females (A) and males (B) showed significant reduction in discrimination ratio scores compared to chance (0.5) in the Novel Object Recognition task. Data are also displayed as percent change from naïve controls, for which both females and males showed significant reductions compared to naïve averages (0%) (C). PME females show a reduction in percent alternation scores in the Spontaneous Alternation task (D), whereas males do not show any significant differences in percent alternation scores across the three exposure groups (E). Data points represent individual animal scores and error bars represent SEM. # p < 0.05 compared to chance, ## p < 0.01 compared to chance, ^ p < 0.05 compared to naïve, *** p < 0.001 as percent change compared to naïve average, & p < 0.05 compared to both naïve and water control groups.

Discrimination ratio data have also been shown as percent change compared to naïve controls (**Fig. 7C**). PME males and females showed significant decreases compared to naïve offspring (theoretical mean = 0) (females: *t*(7)=6.058, *M* = −28.78, p=0.0005; males: *t*(7)=5.537, *M* = −33.67, *p*=0.0009), while water-exposed offspring did not significantly differ compared to naïve offspring (females: *t*(9)=1.243, *M* = −9.06, *p* =0.245; males: *t*(9)=0.209, *M* = −2.53, *p*=0.839).

To ensure results were due to differences in encoding or object side preference during the familiarization day, time spent investigating the objects on familiarization day was also scored. Neither sex showed any significant exposure differences in time spent investigating objects (females: *F*(2,23)=0.522, *p*=0.600; males: *F*(2,24)=0.823, *p*=0.451) or preference for either object during the 10 minute familiarization period (females: *H*(2)=0.5575; *p*=0.750; males: *F*(2,23)=0.255; *p*=0.777) (**Supplementary Fig. 2**).

### 3.6. PME impairs female working spatial memory in the Spontaneous Alternation task

Finally, we examined the effect of PME on adult working spatial memory with the Spontaneous Alternation task. PME females showed a significant reduction in percent alternation scores (*F*(2,21)=5.272, *p*=0.014), with post hoc revealing significant reduction in PME females compared to both water (*p*=0.027) and naïve (*p*=0.027) controls (**Fig. 7D**). Interestingly, males did not show any significant differences in percent alternation scores across the three groups (*F*(2,21)=0.139, *p*=0.871) (**Fig. 7E**). It is important to note that we did discover a significant increase in number of arm entries overall in the water-exposed males (*F*(2,21)=3.977, *p*=0.034), with post-hoc revealing water-exposed males made significantly more arm entries compared to PME males (*p*=0.043) **(Supplementary Fig. 3)**. In order to correct for this, we rescored the male percent alternation scores only up to the average number of arm entries in the naïve group (38); however, this did not change the findings (*F*(2,21)=0.809, *p*=0.457). Females did not show any significant differences in the total number of arm entries (*F*(2,21)=0.104, *p*=0.902).

## 4. Discussion

The present study aimed to begin characterizing the long-term memory impairments following PME, including assessment of hippocampal DGC physiology in adult male and female offspring. We have shown that PME leads to greater inhibition and decreased excitability of DGC only in female offspring. The electrophysiological changes are accompanied by impairments in female working spatial memory and recognition memory in both sexes. These findings support previous studies showing impairments in recognition memory following PME in young male and female rodents^8–10^ and impairments only in adolescent female working memory^9^. Importantly, our data suggest that both effects persist into adulthood.

Very few studies have explored neural disruptions in adult offspring following PME, and here we show that DGC function is disrupted within the hippocampus of adult female offspring. DGCs from methadone-exposed females required more current to elicit an AP, indicating decreased excitability. However, this effect was abolished when blocking GABA_A_ receptors, suggestive of alterations in the inhibitory GABA system. In support of this, sIPSC amplitude was increased in the PME females, indicating increased inhibition of DGCs in female offspring. We did not find any significant changes to sIPSC frequency, suggesting a postsynaptic alteration in GABA transmission, such as changes in receptor expression or function. Previous work has shown alterations in the hippocampal GABA system, with a pronounced decrease in GAD67 and parvalbumin protein expression particularly in female adolescents following PME^9^. Additional work has shown impaired synaptic plasticity and decreased GABAergic neurons in the DG of juveniles^11^ and impaired hippocampal synaptic plasticity in adult rats^18^ following prenatal morphine exposures. These studies, in combination with our current findings, suggest a potential developmental shift from decreased function of the GABA system early in life to an increase in inhibition in adulthood following PME, potentially through compensatory mechanisms. Further experiments are needed to clarify these developmental changes and more closely examine GABA function within the hippocampus following prenatal opioid exposures (POE). Interestingly, male PME offspring showed an increase in time to first action potential when all synaptic activity was blocked, indicative of some impairment in excitability. Nonetheless, our data suggest that DGCs in females may be particularly sensitive to the long-term effects of PME.

Despite females exhibiting more alterations in DGC function, we found similar impairments in recognition memory between males and females. However, only females displayed deficits in working spatial memory. While both NOR and Spontaneous Alternation are hippocampal-dependent^31,32^, it is important to note that the Spontaneous Alternation task is not as dependent on the DG^31^ as the NOR task^22,33^. Additionally, object recognition and spatial memory tasks have been shown to be heavily dependent on adult hippocampal neurogenesis^22,34^, making this a strong candidate for further identification of mechanisms disrupted by this exposure that may coincide to produce the identified behavioral deficits. In support of neurogenesis as a potential mechanism, PME adolescents show reduced mature BDNF^9^, a trophic factor that is critical in regulating neurogenesis.

The sex-dependent nature of the effects of prenatal opioid exposure (POE) in general has been a major focus, and our findings support the need for further examination of both sexes following PME. Importantly, much of the clinical literature relating to POE outcomes in newborns and young children suggest boys may be more vulnerable to the developmental and cognitive outcomes of these exposures^35–38^. However, very little research has followed-up on these effects as children progress into adulthood, especially with PME. Preclinical literature has also shown sex-dependent effects following POE, including higher postnatal death rates in male pups following PME^39^, sex-dependent alterations to the rewarding properties and locomotor effects of alcohol in PME adolescents^40^, and higher vulnerability for hippocampal BDNF dysfunction in prenatal morphine-exposed adolescent and adult females^12^,^13^. This underscores the need for further evaluation of the sex-dependent nature of effects across the lifespan to identify what changes may emerge as offspring develop through puberty and into adulthood.

There were several limitations of this study that should be noted. First, the naïve animals in the NOR task did not acquire the preference for the novel object significantly above chance. Both naïve and water controls showed average discrimination ratios greater than chance, but not statistically significant, whereas PME offspring did show scores significantly below chance. This raises the question of whether this effect is impaired recognition memory or demonstrates neophobia for the novel object in PME offspring. Based on previous studies that do show impaired recognition memory in this task following PME at younger ages^8–10^, combined with evidence for lack of neophobia in this task^41^, and no studies to our knowledge showing neophobia as a result of POE, we hypothesize that our observations are indicative of alterations in recognition memory. However, without further optimization of our testing protocol, the neophobia effect cannot be ruled out.

Additionally, our data reflect potential prenatal stress effects in arm entries during the Spontaneous Alternation task, where the water-exposed males did not resemble the naïve offspring. We feel the inclusion of both control groups is essential in identifying these subtle differences that may be better explained by prenatal stress rather than PME. However, the stress during this exposure does add to the translatability of our model, as most women on methadone maintenance treatments will experience substantial stress during this period^42,43^.

Alternatively, we also acknowledge the translational limitations of this PME model, as the dams do not have a history of opioid exposure nor are they dependent on an opioid before pregnancy.

Finally, in our electrophysiology excitability measures, we standardized all cells by holding them at −70mV to control for RMP variability when conducting the excitability and AP firing experiments. While this may have produced a floor effect and masked potential group differences in AP firing, a significant difference in rheobase was still detectable in females. Nonetheless, future experiments should examine these measures of neuronal excitability at both −70mV and RMP.

The present study provides novel information about the persistent nature of deficits in PME offspring. This is critical in continuing to identify the impact of methadone use during pregnancy and can guide future research toward effective preparation for exposed individuals as they age into adulthood, particularly as the largest cohort of exposed individuals arising from the current opioid crisis have yet to reach adulthood. We have identified long-term neurobiological impairments in exposed offspring, and plan to continue assessing function of the adult hippocampus, as well as other cognitive behaviors that may be affected long-term.

Finally, the sex-dependent nature of effects from POE has been an interesting phenomenon for several decades^35,36,38,44^, and current studies point to similar sex-dependent characteristics following methadone exposures^9,39,40^. This calls for further examination and continued assessment in both sexes to clarify dynamic effects throughout the lifetime. Taken together, this study adds to current literature showing substantial deficits from PME, ultimately lasting into adulthood. This information should be taken into consideration as this medication remains a standard treatment for OUD during pregnancy and may guide future studies to prepare for proper support strategies for methadone-exposed children as they develop.

## Acknowledgements

We would like to thank Allison Regan for her assistance in exposures and NOR data collection and Dana Silberstein for her assistance in PME monitoring. We would also like to thank Lisa Savage and Nicole Reitz for their assistance in behavioral protocols and data consultation. This work was funded by the Binghamton University Psychology Department and Harpur Faculty Research Grant.

## Author Contributions

MEG, RM, and MRD were responsible for the study concept and design. MEG and RM contributed to the acquisition of behavioral data. MEG collected electrophysiological data. MEG, RM, and MRD conducted data analysis and interpretation of findings. MEG drafted the manuscript. MEG and MRD reviewed and edited manuscript. MRD was responsible for funding acquisition and project administration. All authors critically reviewed content and approved final version for publication.

## Data Availability

The data that support the findings of this study are available from the corresponding author upon reasonable request.

## Notes

### Competing Interest Statement

The authors have declared no competing interest.

