## Supplementary Figures for "Prenatal methadone exposure leads to long-term memory impairments and disruptions of dentate granule cell function in a sex-dependent manner"

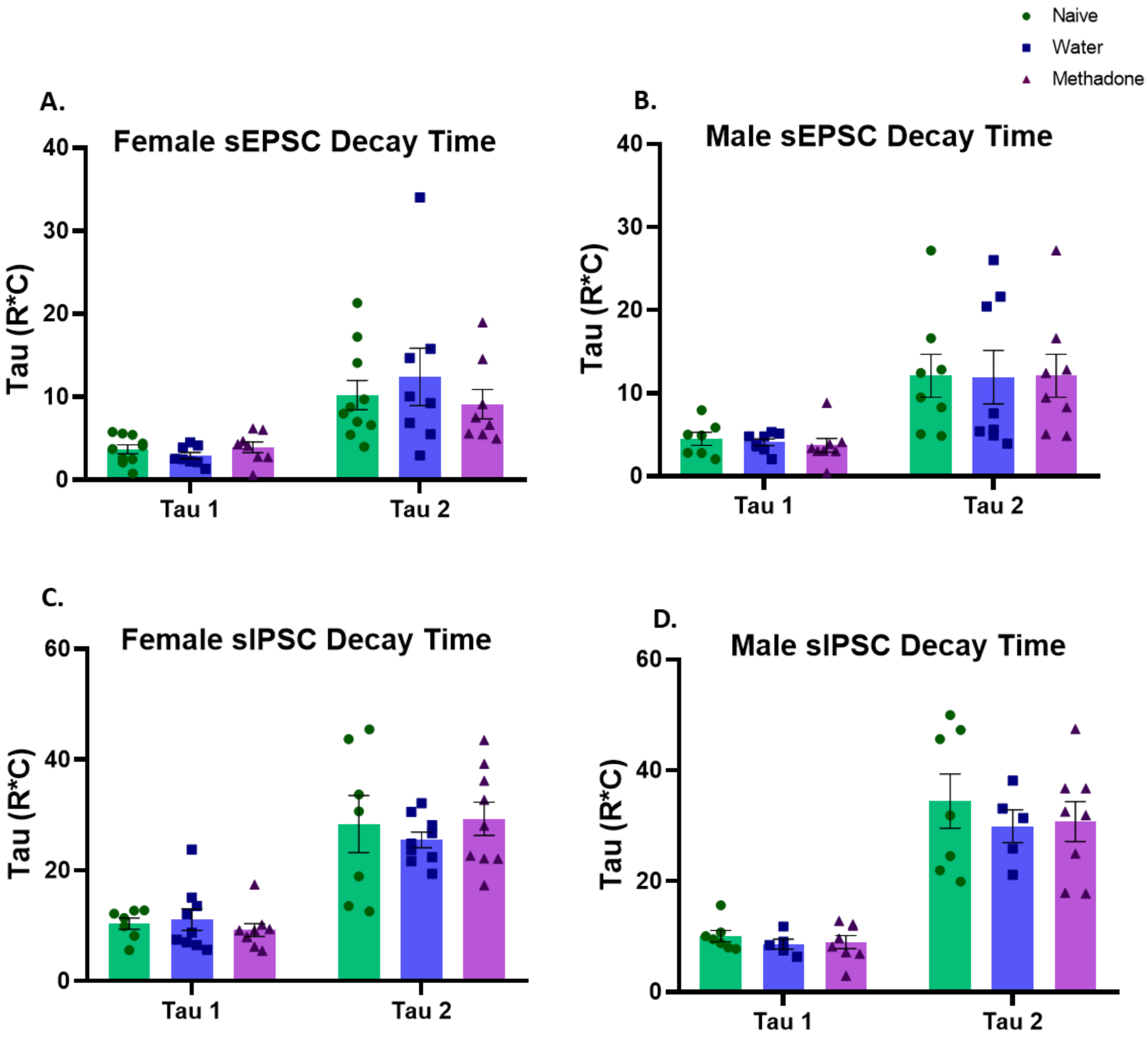


**Supplementary Figure 1.** PME does not alter decay time for male or female sEPSCs or sIPSCs. Female (A) and male (B) offspring show no significant differences in sEPSC decay time across the three groups. Additionally, neither females (C) or males (D) show significant differences in sIPSC decay time across the three groups. Data represent individual cells, with up to three cells per animal. Error bars represent SEM.

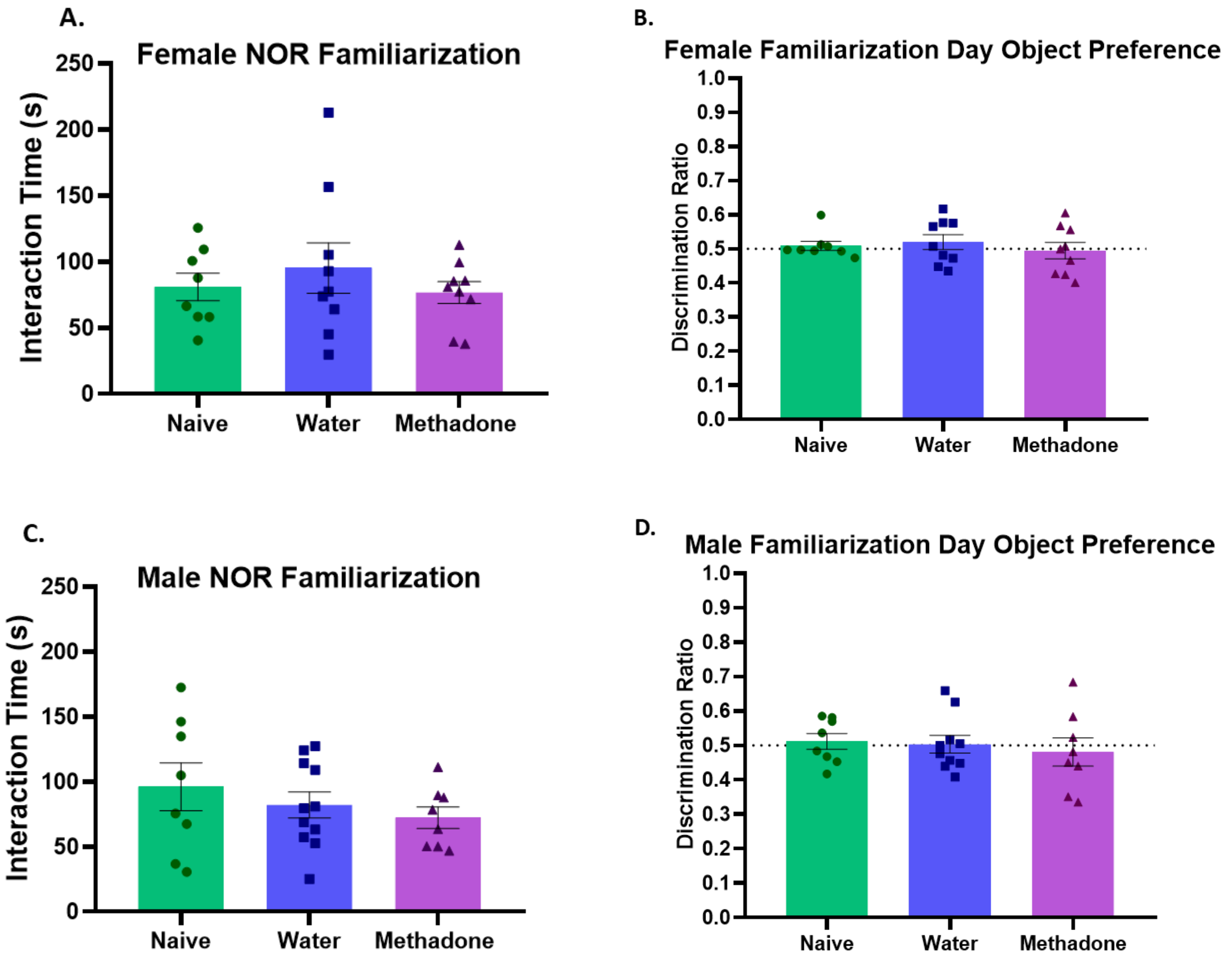


**Supplementary Figure 2.** Results showed no significant differences in the total time spent investigating objects (encoding) and no side preference when investigating the identical objects on the familiarization day of the Novel Object Recognition for females (A-B) or males (C-D). Preference data are displayed as discrimination ratios for the left side object (left object investigation / total investigation of both objects). These data support our results that it is a PME-induced effect on recognition memory during the test day. Data points represent individual animals scores; error bars represent SEM.

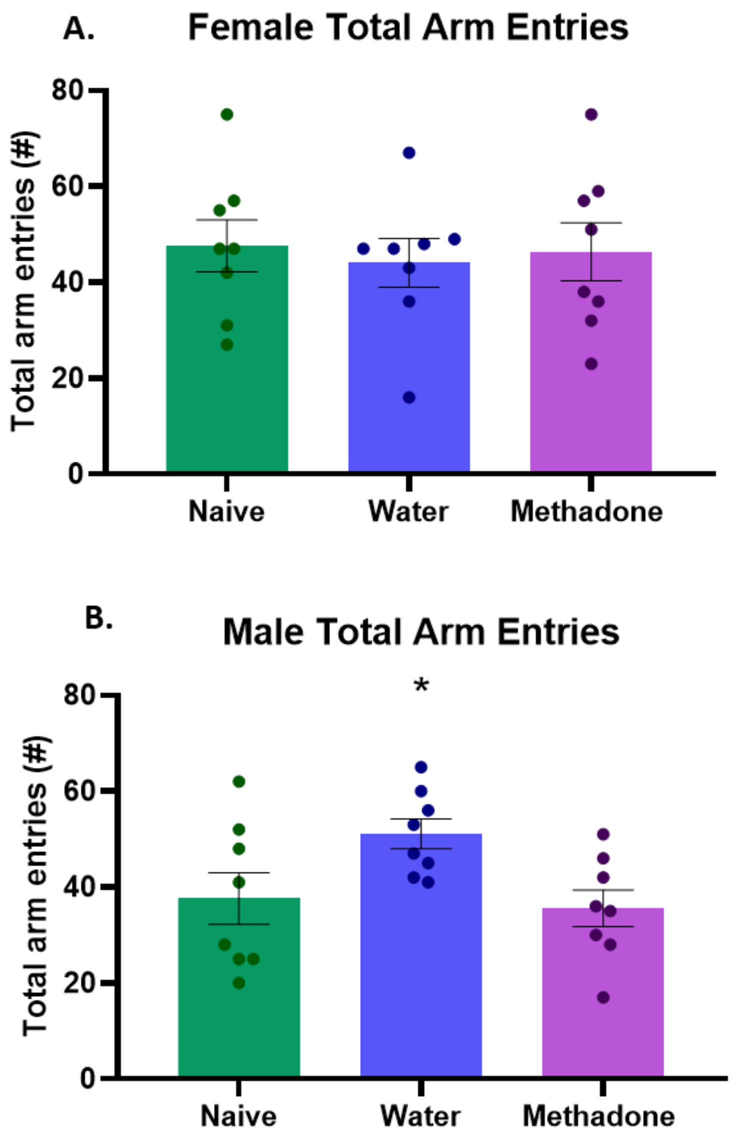


**Supplementary Figure 3.** Female offspring did not show any significant differences across the three exposure groups (A). Water-exposed male offspring showed significantly more arm entries in the Spontaneous Alternation task (B). Data points represent individual animal scores; error bars represent SEM. * $p < 0.05$ compared to PME animals.
